# A Novel Magnetic Field Device: Effects of Magnetic Fields on Planktonic Yeasts and Fungal Mats

**DOI:** 10.1101/2024.04.09.588774

**Authors:** Akila Bandara, Enoki Li, Daniel A. Charlebois

## Abstract

Microorganisms evolved within the geomagnetic field and can be affected by magnetic field exposure. However, the mechanisms underlying many magnetic phenomena in microbes remain to be elucidated. We develop a 3D-printed magnetic field exposure device to perform experiments on microbes. This device is designed in AutoCAD, modeled in COMSOL, and validated using a Gaussmeter. Using the magnetic field exposure device, we perform static magnetic field experiments on different strains of the budding yeast *Saccharomyces cerevisiae*. We find that static magnetic field exposure slows the spatially-structured expansion of yeast mats that expands in two dimensions, but not yeast mats that expand in three dimensions, across the surface of semi-solid media. We also find that magnetic fields do not affect the growth of yeast cells in well-mixed liquid media. This study provides a novel device for magnetic field exposure experiments on microorganisms and advances our understanding of the effects of magnetic fields on fungi.

**Why it matters:** Microorganisms have evolved to function, survive, and reproduce in Earth’s magnetic field. However, the mechanisms underlying magnetic phenomena in microorganisms are unknown. This is especially true for fungi, which are important microorganisms for microbiological research, industrial application, and infectious disease. To elucidate mechanisms driving magnetic phenomena, we need devices to perform controlled experiments in a variety of conditions. We develop a 3D-printed magnetic field exposure device using computer-aided design, physics modeling software, and a magnetometer. Using this novel magnetic field device, we discover that magnetic fields can slow the growth of yeast on agar plates, but that magnetic fields do not affect the growth of yeast in liquid media.

## 1. Introduction

Electromagnetic fields (EMFs) shape the adaptation and evolution of life on Earth [1]. Living organisms including sharks, bees, and birds use EMFs to sense, navigate, and migrate [2]. Magnetic fields (MFs) have been shown to affect the germination of plants [3] and the orientation of blood cells [4, 5]. Magnetotactic bacteria can align with external MFs by biomineralizing magnetic nanoparticles (magnetite or greigite) inside organelles called magnetosomes, which is thought to aid bacteria to reach regions of optimal oxygen concentration [6]. Magnetic nanoparticles have been implemented in cell labeling and imaging [7], as well as targeted drug delivery applications [8]. Despite the advantages of EMFs for living microorganisms, it is important to investigate the detrimental effects of EMF exposure [9].

Due to a short replication time, ease of culture, and well-characterized eukaryotic genetic background, the budding yeast (*Saccharomyces cerevisiae*) is a ubiquitous model organism in molecular biology [10]. *S. cerevisiae* has been used as a model organism in EMF and MF exposure studies [11, 12, 13, 14, 15, 16]. For instance, the orientation of individual *S. cerevisiae* was found to align with a static MF during budding [11]. Exposure to a strong 50000 *G* vertical MF resulted in changes in the sedimentation pattern of yeast cells depending on their location in the culture dishes [13]. However, in the same study, no changes in gene expression were observed after exposure to a 50,000 *G* MF (for 2 and 24 hours) or to a 100,000 *G* MF (for 1 hour).

Previous studies highlight the necessity to study the isolated effects of MFs on fungi. For example, exposing plant pathogenic fungi to low-frequency EMFs and MFs separately yields diverging results: EMFs (1 *G* at 50 *Hz*) were found to have no effect on the growth of mycorrhizal fungi [17], whereas MFs (1 *−* 10 *G*) slowed the growth of phytopathogenic fungi [16]. Another study found that exposing non-pathogenic *S. cerevisiae* cells to stronger EMFs (50 *Hz* with inductions up to 10 *mT*, with exposure times up to 24 minutes) decreased the viability of yeasts cells and slowed their growth [18]. More recently, a lattice-based Monte Carlo simulation framework was developed to investigate the effects of nutrient concentration and MF exposure on yeast colony growth and morphology [19]. Simulation of this framework predicted that prolonged MF exposure will decrease colony growth and alter colony morphology in a nutrient- and ploidy-dependent manner. Despite this research, our knowledge of the effects of MFs on fungi remains limited [16]. Furthermore, though there are many studies on organisms exposed to EMFs (e.g., [18, 20, 21, 22, 23]), the effects of static MF on the growth and development of microorganisms largely remain to be investigated [11].

Yeast exist as part of microbial communities, including colonies, biofilms [24] and “mats” [25, 26, 27, 28, 29]. A yeast mat is a morphologically complex, colony-like structure that requires the expression of the flocculin gene *flo11*, which encodes the *Flo11* surface adhesion protein. Chen *et al.* [30] cultured strains of genetically engineered *S. cerevisiae* cells with and without the *flo11* gene, respectively called TBR1 and TBR5. “Wild-type” TBR1 mats (formed from TBR1 cells with a functional copy of *flo11*) displayed a rough surface, pattern forming phenotype, whereas mutant TBR5 mats (formed from cells lacking a functional copy of *flo11*) displayed a smooth surface phenotype. TBR1 mats grew faster and larger compared to TBR5 mats on semi-solid agar plates; TBR1 also outcompeted TBR5 in competition assays. TBR1 mats are constrained to expand in two dimensions along the agar surface due to the expression of *Flo11* in TBR1 cells. As a result, TBR1 mats had a higher growth rate (fitness) than TBR5 mats, which expand in three dimensions, resulting in a slower mat expansion rate and size along the agar surface. In contrast, there was no difference in fitness between TBR1 and TBR5 strains in liquid culture experiments [30]. Other studies have found similar results for TBR1 and TBR5 growth rates in semi-solid and liquid media cultures [25, 31].

In this study, we develop a MF device to examine the effects of MFs on microbes due to extended periods (30 to 40 days) of exposure to a static MF in the range of 350 *G* to 1500 *G*. The device consists of two Neodymium (Nd_2_Fe_14_B) magnets that exposes multiple biological replicates to a MF; the device can be placed inside of an controlled environment for the duration of experiments. We design the MF exposure device in AutoCAD [32] and model it using COMSOL Multiphysics [33]. We then 3D print the optimized device and measure the MF inside the exposure region, which is compared to numerical COMSOL simulations. To highlight the utility of our device, we perform magnetic field experiments on yeast cultures inside of an environmental chamber. We find that 1) MFs slow the growth of TBR1 mats on semi-solid agar media, but MFs do not affect the growth of TBR5 mats; 2) TBR1 and TBR5 mats increase their growth rates on agar media in the presence and absence of MFs; and 3) MFs do not affect the growth rate of planktonic TBR1 and TBR5 cells in well-mixed liquid media.

## 2. Materials and Methods

### 2.1. Fabricating the MF exposure device

To create a homogeneous MF to expose biological samples of microbes growing in cell culture tubes or Petri dishes, we used two N52 grade Nd_2_Fe_14_B block magnets (Amazing Magnets Inc, Catalog no. Q500Y-N52). The device’s ability to hold multiple replicates (five Petri dishes or nineteen culture tubes) for each exposure experiment is essential to ensure the repeatability of the results and to test for statistical significance. The horizontal and vertical configurations of the block magnets in the device permit MF exposure emanating from two different directions (Figs. 1A,B and C1). With dimensions of 101.6 *×* 101.6 *×* 12.7 *mm*^3^, these magnets were capable of producing a magnetic flux density (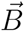) of approximately 14400 *G* at the magnets surface, which is five orders of magnitude stronger than the Earth’s MF (0.24 *G* to 0.66 *G* depending on the latitude [34]).

**Figure 1:**
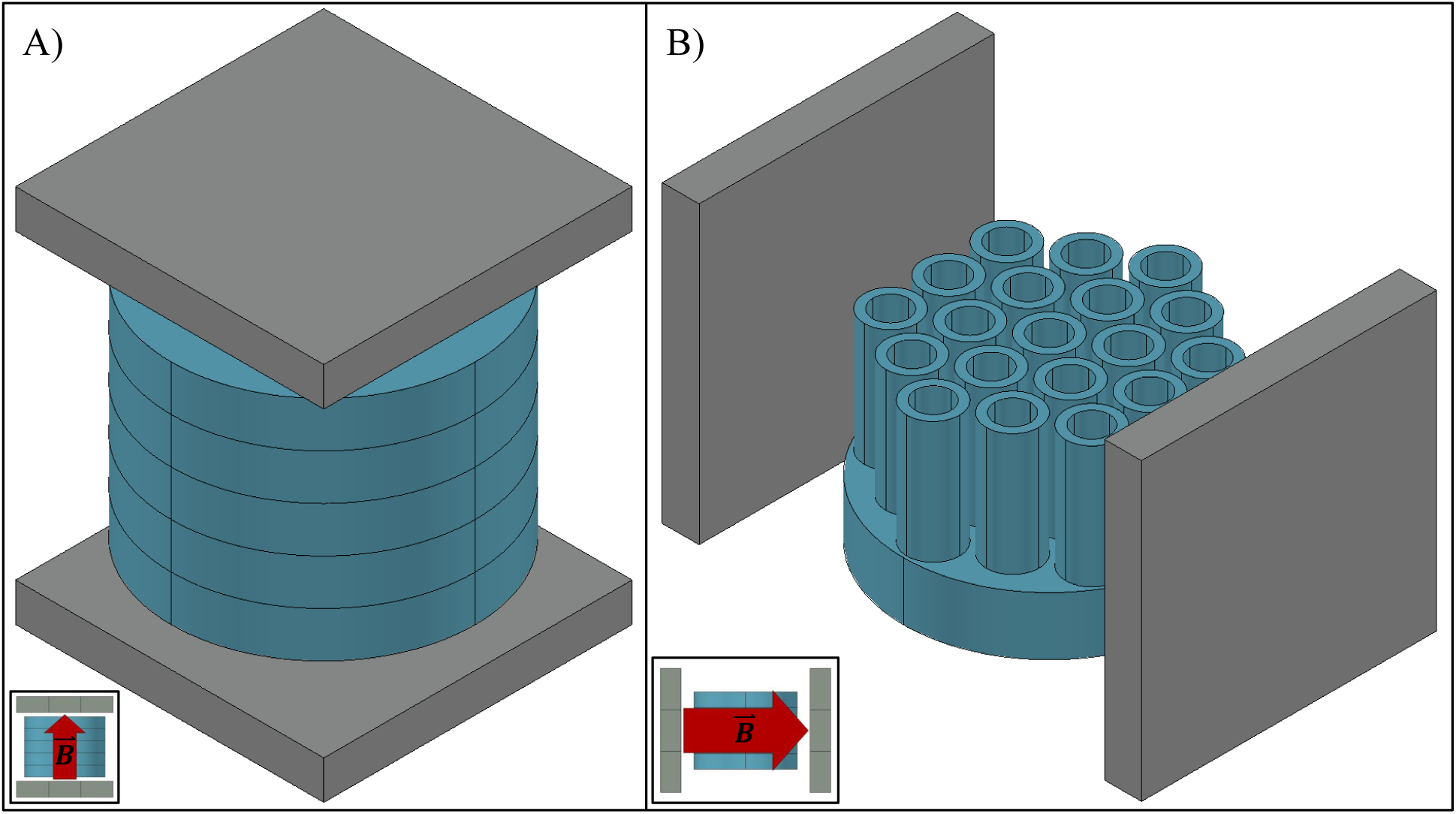
AutoCAD [32] schematics showing the different configurations of the magnetic field device. (A) Schematic of the vertical configuration of the device, which was can be used to expose yeast grown on semi-solid media in Petri dishes to a static vertical magnetic field (MF). Petri dishes are shown in blue. (B) Schematic of the horizontal configuration of the device, which was used to expose yeast grown in liquid media in cell culture tubes to a static horizontal MF. Culture tubes and holder are shown in blue. The grey blocks in (A) and (B) represent the Neodymium magnets; insets show the direction of the magnetic flux density (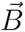).

The MF exposure device materials were selected to minimize interaction between the magnets and the scaffolding. Considering that microorganisms will typically be grown on semi-solid media in Petri dishes or in liquid media in culture tubes (see Section 2.3), it was necessary to fabricate the MF device using a material that had compatible magnetic permeability (*µ*) to the Petri dishes and culture tubes. This ensures that the homogeneous MFs produced by the block magnets would have the least possible interference during its path towards the exposure samples [35]. Since two strong magnets are in close proximity (132 *mm*), the device materials need to be sufficiently strong to withstand a pull force of 5.78 *N* without compromising the structural integrity of the device. Furthermore, as the device is intended to perform experiments on biological samples, it was designed to be disassembled so that the components can be cleaned and sterilized.

We used AutoCAD [32] to design a MF device with two interchangeable configurations: a horizontal 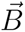 configuration and a vertical 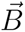 configuration (see Fig. C1). COMSOL Multiphysics [33] was used to simulate the MF inside the device to optimize the design (see Section 3.1). The “Magnetic Fields, No currents” interface in the AC/DC module of COMSOL was used to simulate the 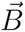 values within exposure region of the MF device via a finite element method [36]. The MF strength (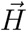) and the 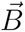 resulting from the permanent magnets in the exposure device are respectively described by Maxwell’s equations for a static MF [37]:

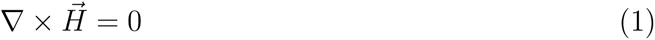

and

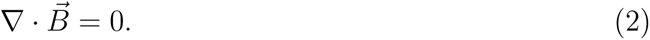

From Equation (1), we can define the magnetic scalar potential (*V_m_*) as:

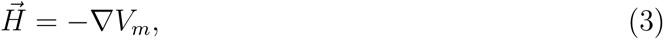

and considering the relation between the 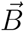 and 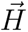 yields:

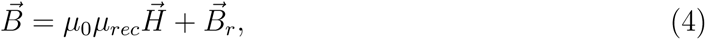

where *µ*_0_, *µ_r_*, and 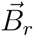 represent the permeability of free space, recoil permeability, and the remnant flux density of the permanent magnet, respectively. By combining Equations (3) and (4) into (2), we obtain:

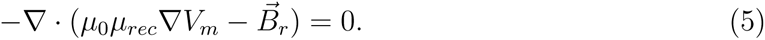

Equation (5) was used in our numerical COMSOL simulations to describe the material properties of the Nd_2_Fe_14_B block magnets. The *µ* of polylactic acid (PLA) was also incorporated into the COMSOL simulations to specify the device material.

We used PLA (RepRap Warehouse, Catalog No. R00100002), a thermoplastic polyester with a low melting point, high strength, low thermal expansion, and with a *µ* comparable to the Petri dishes/culture tubes, provided the necessary durability and versatility to our MF exposure device. The components of the device were 3D printed using PLA filaments with diameters of 1.75 *mm* with Prusa i3 MK3S and i3 MK3S+ printers.

### 2.2. Exposure device MF simulations and measurements

COMSOL multiphysics [33] was used to simulate the MFs of the prototype devices to optimize their design before they were 3D printed. The AutoCAD [32] designs of the device were imported into COMSOL and the properties of each material used in the device were defined based on their value of *µ* (see Table A1).

The 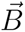 in the exposure region of the MF device was experimentally measured using a Gaussmeter (Alphalab Inc., Model GM2 Gaussmeter), which operates based on the Hall effect [38]. To create a 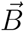 map, a custom cylindrical Gaussmeter probe holder containing 83 equally spaced rectangular holes was designed and 3D printed (Fig. C2A). This permitted us to measure the 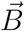 in each layer at specific positions inside the exposure chamber (Fig. C2B). We were able to limit the 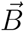 mapping to three layers due to the symmetry of the MF. Four 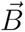 readings were obtained for each position in the these layers. The average of these 4 readings was used to obtain the final mapping of 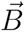.

### 2.3. Yeast MF exposure experiments

Haploid *S. cerevisiae* TBR1 (Σ1278b, mat*α*, *flo11*, tryp) and TBR5 (Σ1278b, mat*α*, *flo11* Δ, tryp) strains were used for the MF exposure experiments [30]. TBR1 (*flo11*) and TBR5 (*flo11* Δ) cells are isogenic apart from the presence or absence of the *flo11* gene, respectively.

Biological replicates of TBR1 and TBR5 were cultured from isogenic colonies in YPD liquid medium at 30 *^◦^C* and shaken overnight on a CO_2_-resistant shaker (Thermo Scientific, Catalog no. 88-881-103) at 150 rpm inside an environmental chamber (Thermo Scientific, Catalog no. 13-067-066). YPD media was made with 5 *g* of yeast extract (Sigma-Aldrich, Catalog no. Y1625), 10 *g* of bacto peptone (BD, Catalog no. 211677), 38 *mg* of adenine (Sigma-Aldrich, Catalogue no. D16), and 7.5 *g* (or a 1.5 % final concentration) of agar (for agar plates: Fisher Scientific, Catalogue no. BP1423), autoclaved in 450 *ml* of Type 1 water, and supplemented with glucose (Fisher Scientific, Catalog no. D16) to a final concentration of 2 %. Liquid TBR1 and TBR5 cultures were inoculated to 5.5 *×* 10^5^ *cells/ml* in 5 *ml* culture tubes (Fisher Scientific, Catalog no. 22-171-606). Agar TBR1 and TBR5 cultures were inoculated with a 2 *µl* drop of 10^7^ *cells/ml* onto agar medium in a “100 *mm*” (outer diameter is *≈* 92 *mm* and inner diameter is *≈* 88 *mm*) Petri dish (Fisher Brand, Catalog no. FB0875712).

For the agar culture-MF exposure experiments, twenty 100 *mm* Petri dishes (half seeded with TBR1 cells and half seeded with TBR5 cells) were placed inside of four 3D-printed devices. Five TBR1 Petri dishes and five TBR5 Petri dishes were place inside two separate exposure devices (block magnets present), and five TBR1 plates and five TBR5 plates were placed inside two separate control devices (block magnets absent). Considering *flo11*’s function to restrict TBR1 mat expansion along the surface of the agar plate [30], the MF exposure devices were set up in the horizontal configuration (Fig. 1B). The horizontal MF creates an external magnetic force parallel to the plane of expansion of yeast mats. The experimental and control devices were then placed inside an environmental chamber (Thermo Scientific, Catalog no. 13-067-066) and incubated at 30 *^◦^C* and 50 % humidity. Yeast mats seeded with TBR1 or TBR5 cells were grown for 25 days. The yeast mats were photographed daily using a Canon EOS Rebel SL3 camera with a Canon EF-S 3 *mm* f/2.8 Macro IS STM macro lens. The growth rates were evaluated from these photographs as described in Section 2.4.

For the liquid culture-MF exposure experiments, twenty four 5 *ml* culture tubes (Fisher Scientific, Catalog no. 22-171-606; half containing TBR1 cells and half containing TBR5 cells) were placed inside four 3D-printed devices. Six TBR1 tubes and six TBR5 tubes were placed inside two separate experimental exposure devices. Another six TBR1 tubes and six TBR5 tubes were places inside two separate control devices. To maintain consistency with the agar culture-MF exposure experiments, the liquid culture-MF exposure experiments were also set up in the horizontal configuration. The experimental and control devices were then placed inside the environmental chamber on the shaker and incubated at 30 *^◦^C* and 50 % humidity. The liquid TBR1 and TBR5 cultures were grown for 3 days. The growth rates were evaluated and the cultures re-suspended every 12 hours (see Section 2.5).

### 2.4. Agar culture growth rate measurements

Yeast mat area measurements were obtained daily for 25 days to determine the difference between the mat expansion rate of the control and the MF-exposed TBR1 and TBR5 strains. A quantitative analysis of the mat area expansion rates was performed using the image processing software ImageJ [39]. The resolution of the original images was 6000 by 4000 pixels with an aspect ratio of 3:2. To obtain the area expansion rate from the original images, the original images were cropped to a square ratio (1:1) such that the circumference of the Petri dish in the image touched each of the four sides of the image. This enabled the accuracy to be maintained when scaling the image pixels by the length of the Petri dish in each image, as the area evaluation of the mats remained consistent throughout the analysis. After extracting the area of the mats from each image, the area expansion rates were obtained by dividing the total area at the end of each day by the number of days.

### 2.5. Liquid culture growth rate measurements

To determine the difference between the growth rates of the control and MF-exposed TBR1 and TBR5 strains in well-mixed liquid cultures, cells were extracted from TBR1-seeded and TBR5-seeded mats using sterile 20 *µ*l pipette tips (Fisher Scientific, Catalog no. 02-707-432). Specifically, twelve cultures of TBR1 cells were extracted from six TBR1 control and six TBR1 MF exposure plates and twelve cultures of TBR5 cells were extracted from six TBR5 control and six TBR5 MF exposure plates and grown in liquid YPD medium for 3 days. Every 12 hours, cell counts were obtained using an automated cell counter (Corning, Catalog no. C-6749) and the cell cultures were re-suspended to an initial concentration of *N*_0_ = 5.5 *×* 10^5^ *cells/ml* to keep them in log-phase growth. The population growth rate was calculated as follows [40]:

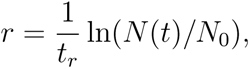

where *t_r_* is the time interval between re-suspensions and *N*(*t*) is the cell count at time *t* after the re-suspension.

## 3. Results

### 3.1. MF Device Optimization and Validation

The MF within the simulated exposure device had a well-defined orientation (Figs. 2A and 3A). Simulations of the vertical configuration of the exposure device produced 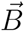 values ranging from 430 to 730 *G* along the horizontal surface of the central Petri dish (Fig. 2B). The 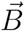 values ranged from 500 to 1400 *G* along the central vertical plane of the exposure chamber (Fig. 2C). Simulations of the horizontal configuration of the exposure device produced 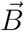 values ranging from 500 to 1400 *G* (Fig. 3B,C). In agreement with theory [35], the simulations predicted negligible interference with the MF when the exposure device is fabricated with materials with compatible *µ* values (Figs. 2A and 3A).

**Figure 2:**
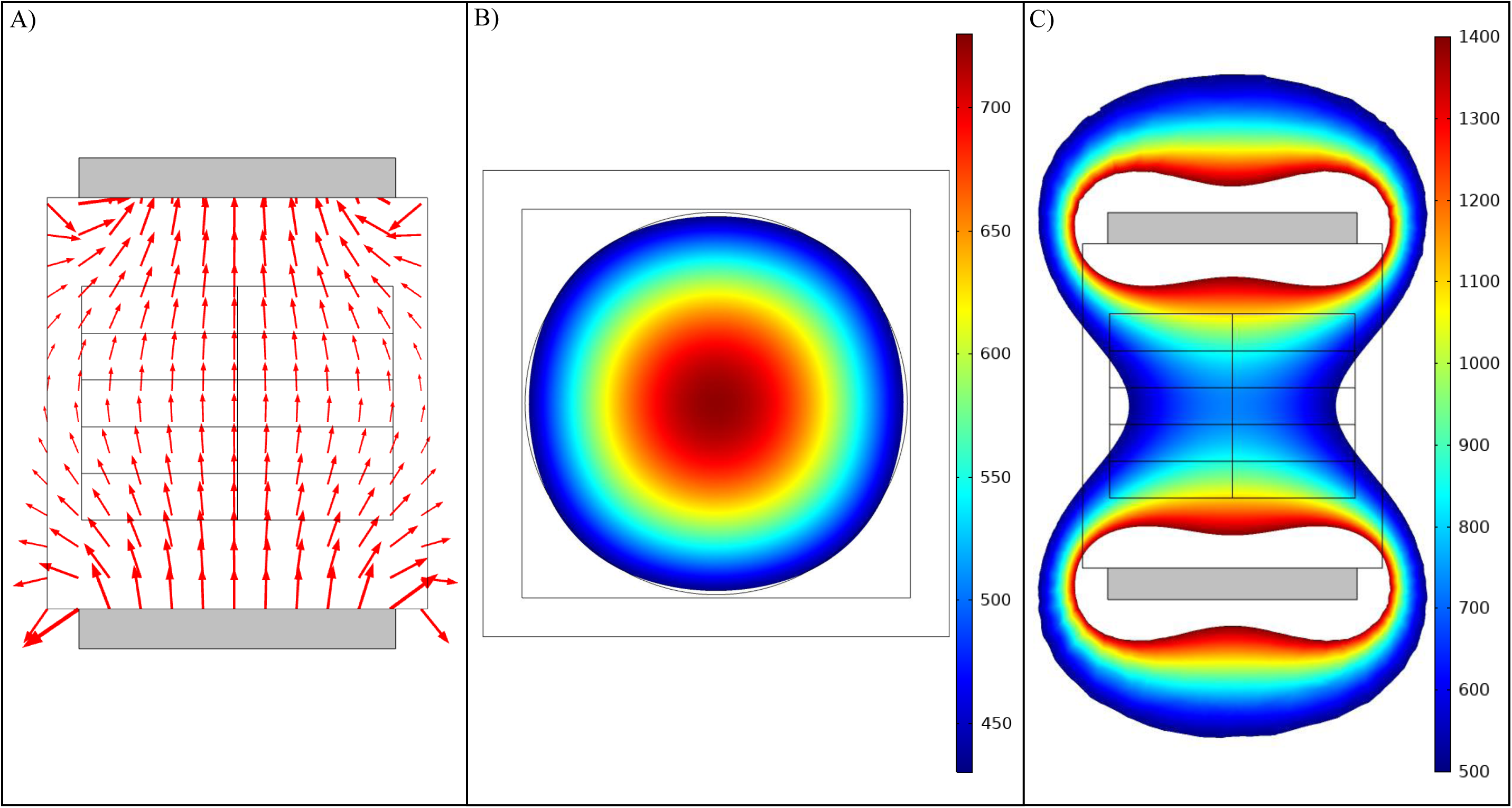
Simulation of the vertical configuration of the magnetic field device. (A) COMSOL [33] simulation of the directional orientation of the magnetic field. (B) COMSOL simulation of the magnetic flux density (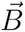) along the central horizontal plane of the exposure device. The point of view of this panel is from above looking down at the central Petri dish. (C) COMSOL simulation of the 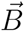 along the central vertical plane of the exposure device. The point of view of this panel is from the side of the exposure device looking at the sides of the Petri dishes. The grey blocks in (A) and (C) denote the Neodymium magnets. The colorbars denote 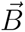 in Gauss (G).

**Figure 3:**
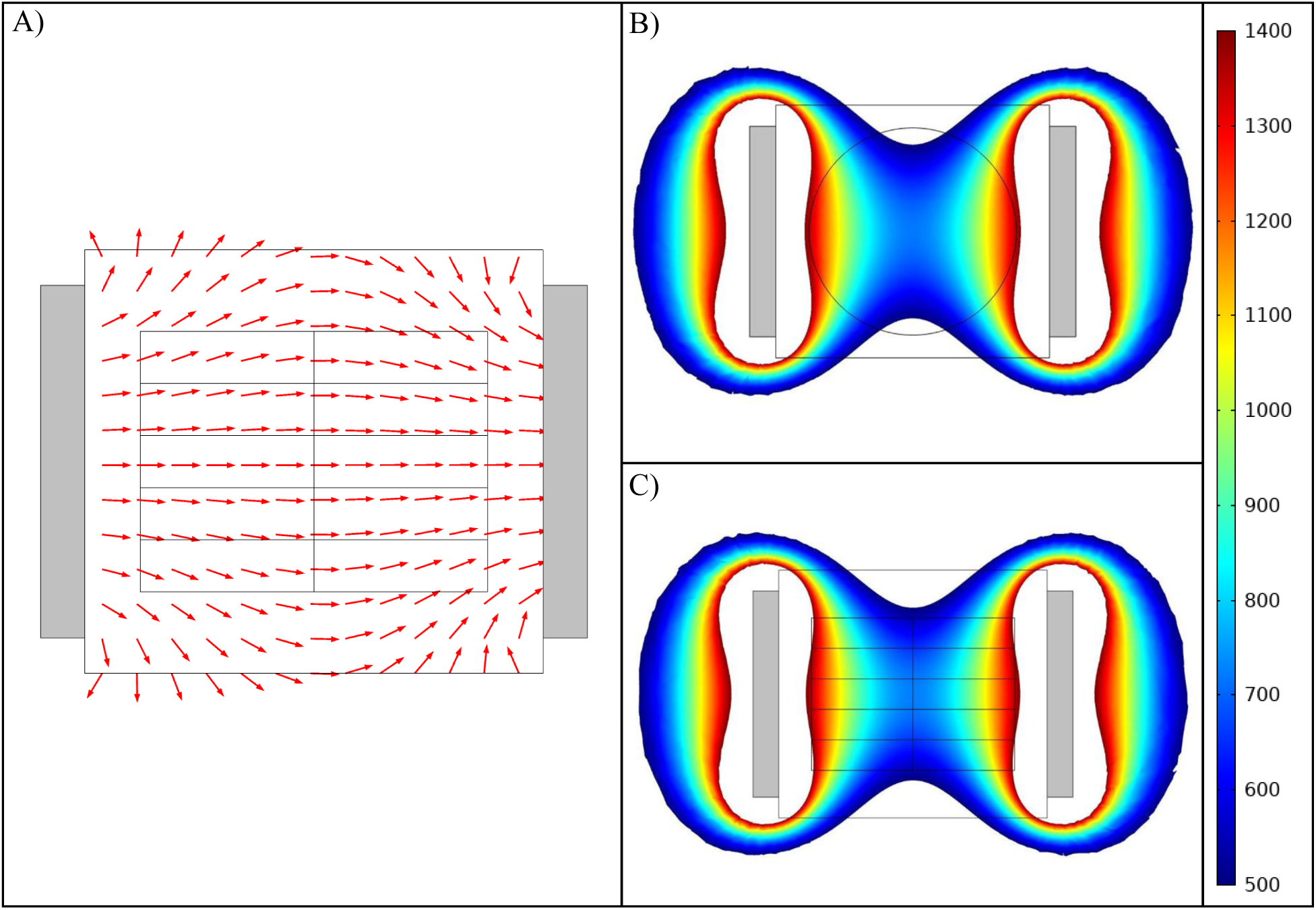
Simulation of the horizontal configuration of the magnetic field device. (A) COMSOL [33] simulation results of the directional orientation of the magnetic field. (B) COMSOL simulation of the magnetic flux density (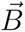) along the central horizontal plane of the exposure device. The point of view of this panel is from above looking down at the central Petri dish. (C) COMSOL simulation of the 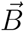 along the central plane from the side view of the exposure device. The grey blocks in (A)-(C) represent the Neodymium magnets. The colorbar denotes 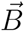 in Gauss (G).

Simulated and experimentally determined 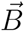 maps for the middle layer (layer 3) of the exposure device are shown in Figure 4. The simulated 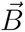 values for layer 3 ranged from 514.6 *G* to 1147.5 *G* (Figs. 4A and C3A). The experimental 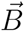 values for layer 3 ranged between 550.4 *G* and 1194.4 *G* (Figs. 4B and C3B). The simulated and experimental 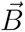 results for layers 1 and 2 are shown in Figures C7 and C8, respectively. On average, each experimental 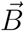 mapping value for layer 3 was 35.5 *G* different than the corresponding COMSOL simulated value (Fig. C4A). This corresponds to a 4.4 % average difference between the experimental and simulated 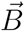 values (Fig. C4B). Results for layers 1 and 2 follow a similar trend for the differences between the experimental and COMSOL simulated 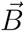 values. For layer 1, each experimental 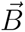 value was 37.2 *G* less than the corresponding COMSOL simulated value on average, which corresponds to a 5.7 % difference between the experimental and simulated 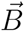 values (Fig. C7). For layer 2, each experimental 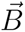 value was 35.0 *G* less than the corresponding COMSOL simulated value on average, which corresponds to a 4.5 % difference between the experimental and simulated 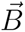 values (Fig. C8). Overall, the experimental 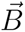 measurements were in good agreement with the simulation results.

**Figure 4:**
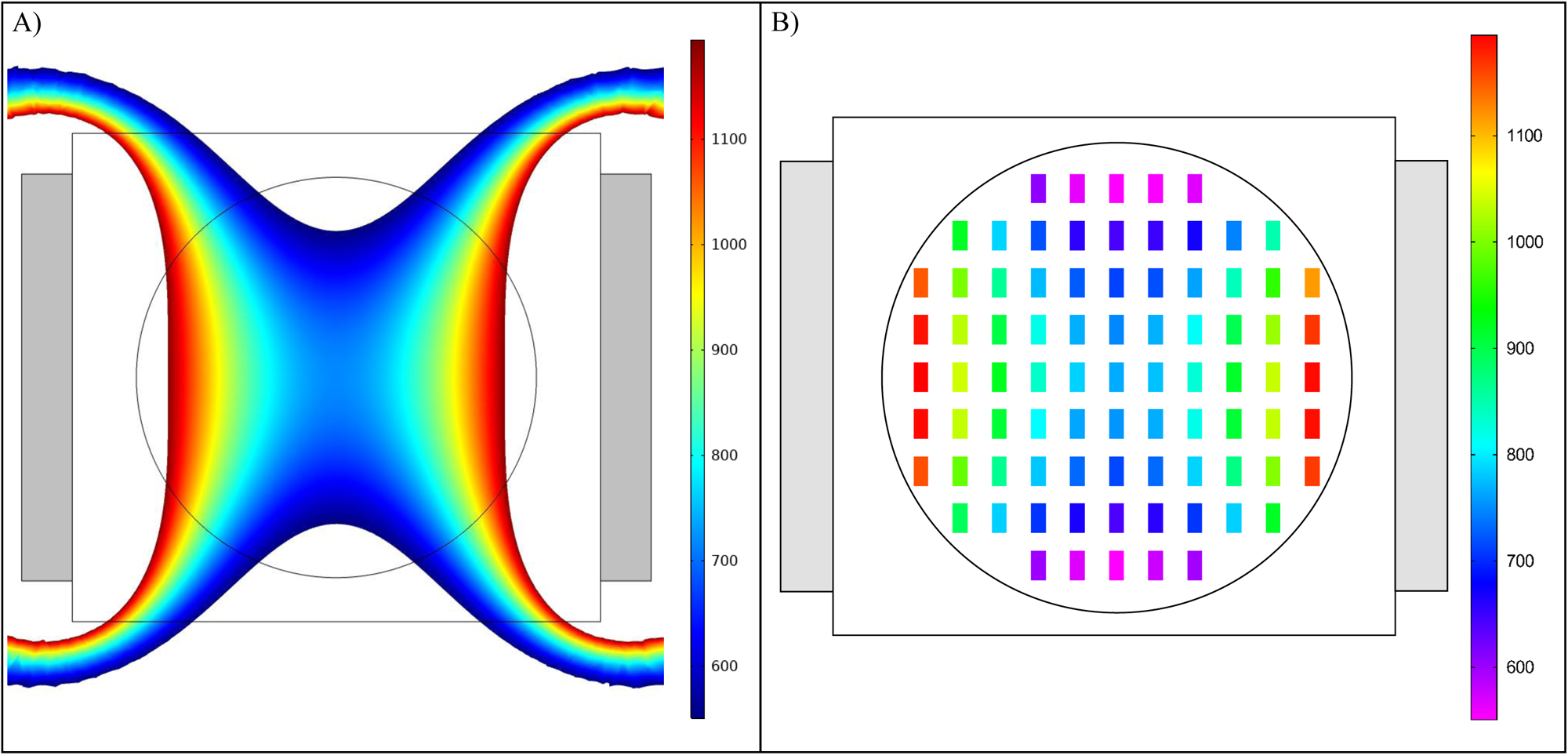
Comparison of simulation and experimental measurements of the horizontal configuration of the magnetic field device. (A) Top view of a COMSOL [33] simulation of the magnetic flux density (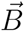) along the middle layer (layer 3) of the exposure chamber of the device. (B) Top view of the experimental 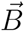 mapping of the device corresponding to (A). The grey blocks in (A) and (B) represent the Neodymium magnets. The colorbars denotes 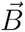 in Gauss (G).

### 3.2. Spatiotemporal yeast mat MF experiments

Magnetic field exposure decreased the TBR1 mat area expansion rate (Fig. 5). Statistically significant differences in the average area expansion rate between the experimental (MF exposed) and control (no MF) groups occurred for days 7 to 22. The average area expansion rate of the control group expanded at an linearly increasing rate (Table B1) until saturation around day 18, and then began to decrease. The saturation and subsequent decrease in the average area expansion rate can be attributed to nutrient depletion as the yeast mats expand across the agar surface [28, 41]. In contrast, the average area expansion rate of the MF exposed group displayed a slower, monotonic and linearly increasing growth rate (Table B1). The average area of control and MF-exposed TBR1 mats expanded exponentially (Table B2).

**Figure 5:**
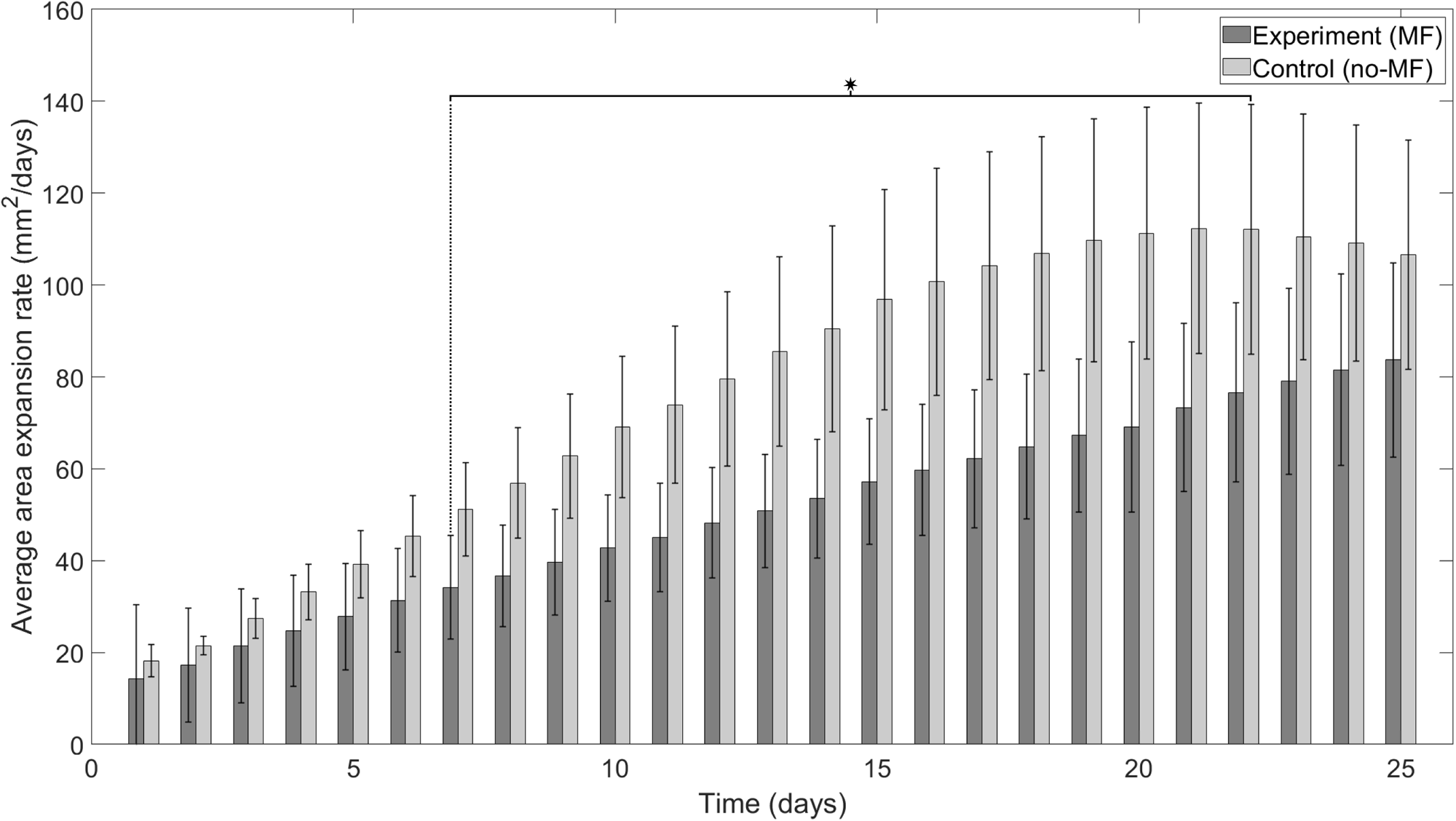
Average area expansion rates of TBR1 yeast mats in agar plates in the presence and absence of a horizontal magnetic field (MF). TBR1-seeded control (no-MF) and experimental (MF) mats grown on agar. An independent samples t-test was performed to compare the average expansion rates for a sample size of N = 5. A significant difference in average area expansion rates independent samples t-test (p < 0.05, denoted by an asterisk) was found for days 7 to 22.

MF exposure did not impact the TBR5 mat area expansion expansion rate (Fig. 6). No significant difference in the average area expansion rate between the experimental (MF exposed) and control (no MF) groups occurred on any day of the experiment. Since the TBR5 expansion rate is slower compared to TBR1 (Fig. 5), the TBR5 average area expansion rates did saturate during the experiment (Fig. 6) as TBR5 mats likely did not sufficiently deplete the nutrients in the agar plates. Experimental and control groups of TBR5 yeast mats displayed a logarithmic average area expansion rates (Table B1), which increased monotonically for the duration of the experiment. The average area of control and MF-exposed TBR5 mats expanded linearly (Table B2).

**Figure 6:**
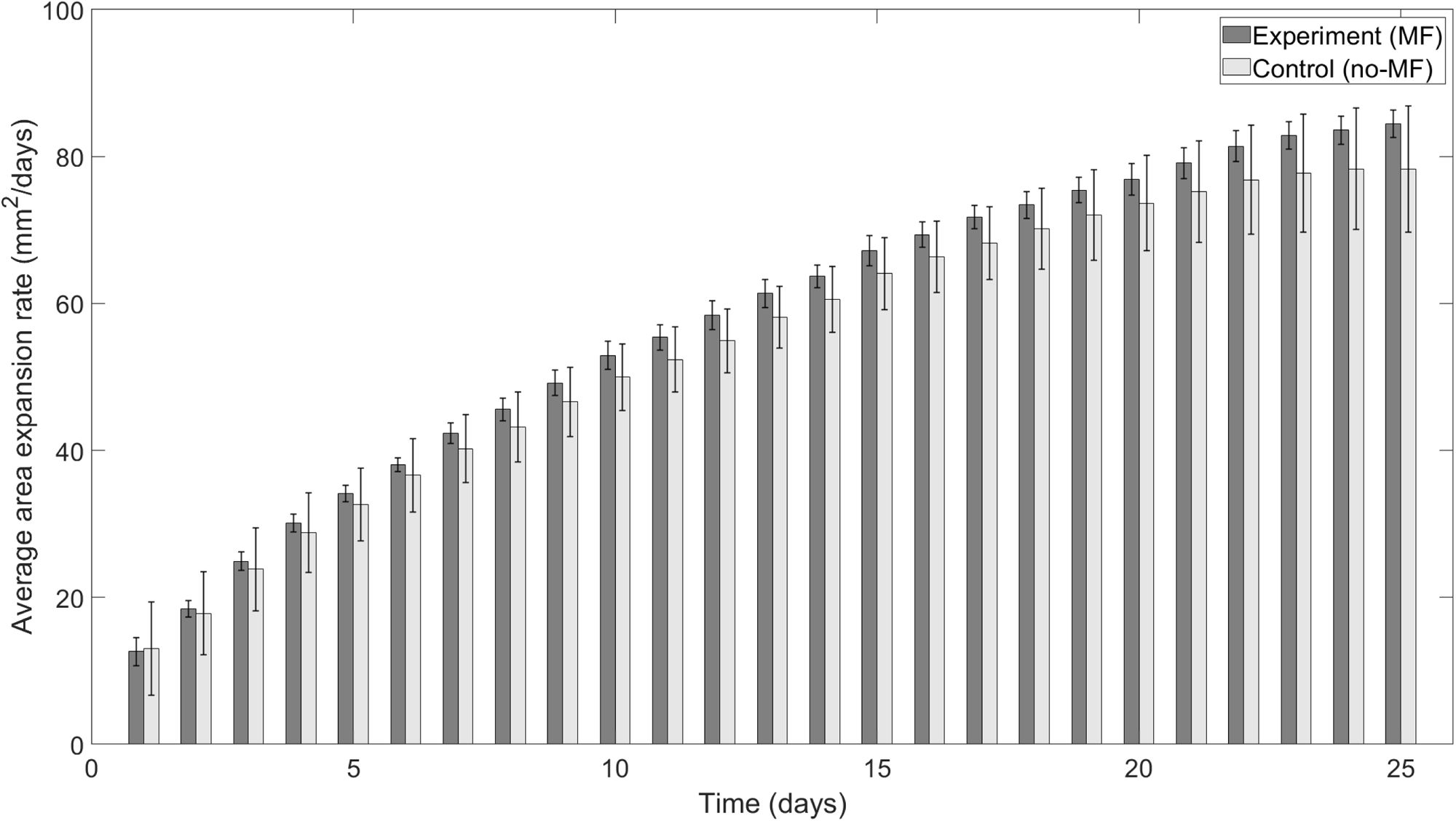
Average area expansion rates of TBR5 yeast mats in the presence and absence of a horizontal magnetic field (MF). TBR5-seeded control (no-MF) and experimental (MF) mats grown on agar plates. An independent samples t-test was performed to compare the average expansion rates for a sample size of N = 3. No significant difference in average area expansion rates (p > 0.05) was found for any day. Note that here N = 3 (not N = 5, as for TBR1) as 2 experimental replicates were discarded due to insufficient agar medium and contamination.

### 3.3. Well-mixed planktonic yeast MF experiments

Overall, MF exposure did not effect the steady-state growth of TBR1 and TBR5 cells cultured in liquid media (Fig. 7). No significant differences were observed in TBR1 average growth rate between the control and MF-exposed at the 12, 36, 48, and 60 hour time points and no significant differences were observed in TBR5 average growth rate between MF exposed and control cultures at the 24, 36, 48 and 60 hour time points. The similarity in the growth rates for TBR1 and TBR5 can be likely be explained by the negation of the function of the *flo11* gene in liquid media. These results are in agreement with previous TBR1 and TBR5 liquid culture experiments [30, 31].

**Figure 7:**
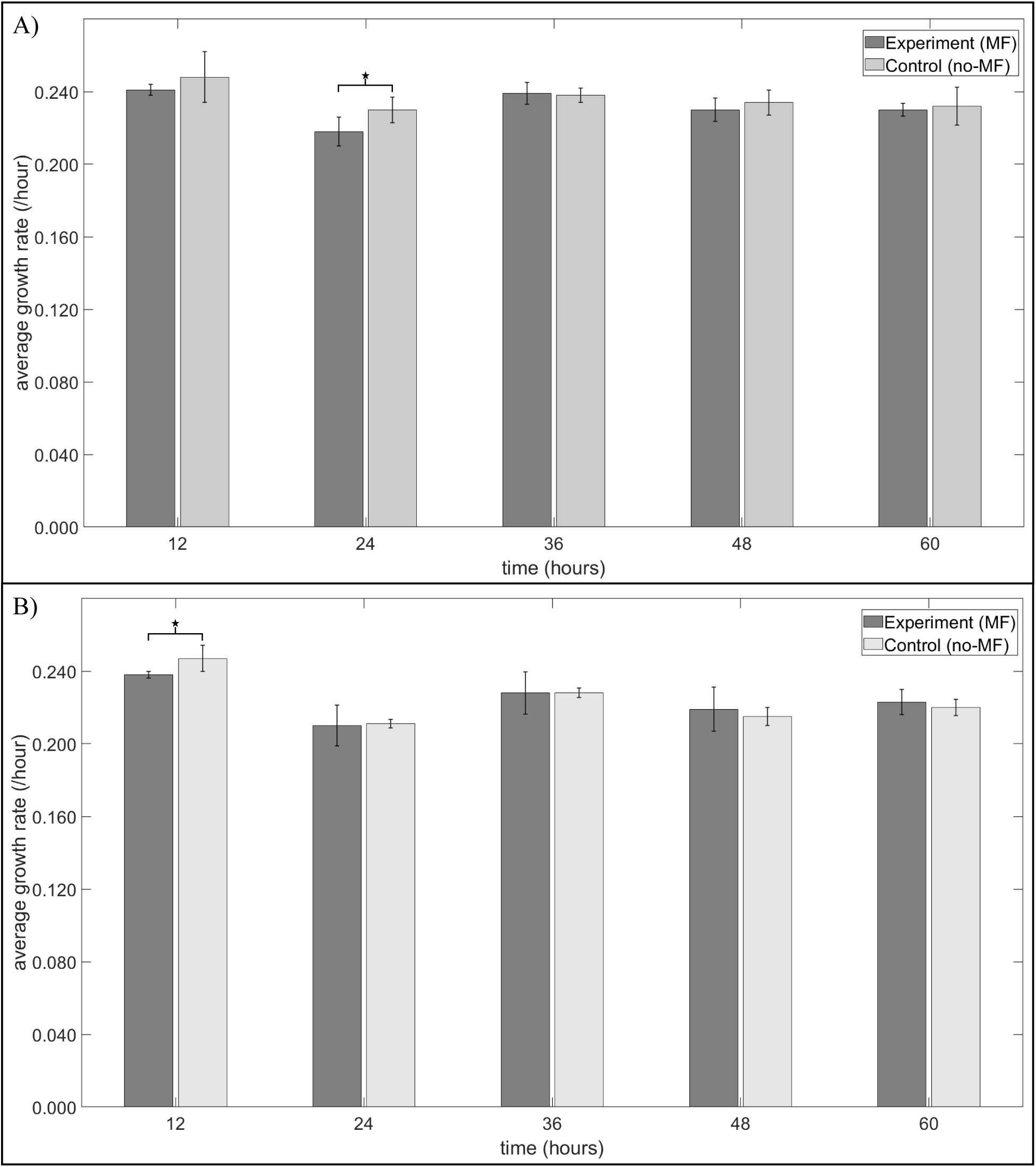
Average growth rates of TBR1 and TBR5 cells in liquid culture in the presence and absence of a horizontal magnetic field (MF). (A) Average growth rates of TBR1. An independent samples t-test was performed to compare the average growth rates between the exposure (MF) and control (no-MF) replicates for a sample size of N = 6. A significant difference (p < 0.05) was only observed at the 24 hour time point. (B) Average growth rate of TBR5. An independent samples t-test was performed to compare the average growth rates between the exposure (MF) and control (no-MF) replicates for a sample size of N = 6. A significant difference(p < 0.05) was only observed at the 12 hour time point.

## 4. Discussion

In this study, we developed a static MF exposure device to perform controlled MF experiments on microorganisms. This 3D-printed device was designed in AutoCAD [32] to have interchangeable horizontal and vertical MF configurations, to hold culture tubes and Petri dishes. The strength, size, and position of the magnets were optimized for MF exposure experiments on the budding yeast *Saccharomyces cerevisiae* using numerical simulations in COMSOL [33] along with Gaussmeter measurements. The AutoCAD, COMSOL, and 3D printing files are freely available for use in other MF experiments (see Supporting Materials). While a previous MF device was designed to investigate the effects of MF on single yeast cells, our MF device provides to ability to study the effects of MFs on communities of yeast cells. By using two large Neodymium magnets, we are able to provide a sufficient volume to generate a homogeneous MF throughout exposure region, which will facilitate MF exposure experiments on yeast mats/biofilms on agar plates and yeast cells in liquid culture. Additionally, while many previous MF exposure experiments were conducted for no longer than 2 days [12, 13, 14, 11], our MF exposure device is able to maintain an uninterrupted MF for longer-term experiments, the importance of which has been previously emphasized [19]. Finally, our device can be placed inside of a standard microbiological incubator or environmental chamber to control for confounding variables, such as temperature, that affect growth and gene expression in yeast [42].

We used our MF exposure device to investigate the effects of MFs on the growth of two *S. cerevisiae* strains in agar and liquid media. We found that MFs slowed the expansion of TBR1 yeast mats on agar plates, but that MFs did not affect the expansion of TBR5 mats. We also found that MFs did not affect the growth of the TBR1 and TBR5 yeast cells in well-mixed liquid media. The decreased expansion rate of MF-exposed TBR1 yeast mats may be attributed to spatial hindrance due the expression of the *flo11* surface adhesion gene in this strain [30] combined with the magnetic properties of microtubules [11, 43, 44]. As the mitotic spindle is composed of [45] and oriented [46] by microtubules, the presence of a horizontal MF may cause the budding yeast cells to align their long axis along the direction of the MF [19, 11]. Partial or complete alignment of cells to the MF at the expanding boundary of a yeast mat may introduce competition among cells attempting to bud into unoccupied space on the agar surface. In TBR1 yeast mats, with the presence of the *Flo11* protein favoring two-dimensional expansion [30], competition introduced by external MF may underlie the significant decrease in area expansion rates of MF-exposed TBR1 mats. MF-induced steric hindrance may also explain the previously observed slowed growth of phytopathogenic fungi exposed to MFs [16]). While MF orientation effects were not found to impact the growth of individual budding yeast cells embedded in agar [11], these effects may have contributed expansion rate of multicelluar yeast mats across the agar surface in our study. While TBR5 cells would experience similar MF orientation effects as TBR1 cells, the fact that TBR5 mats expand in 3D across the agar surface (due to the absence of *flo11* in this strain) may negate spatial hindrance at the expanding mat front. When planktonic TBR1 and TBR5 cells were grown in liquid media, no MF-related effects on growth were observed; this can likely be attributed to the lack of *flo11* functionality in liquid media, thus rendering the TBR1 and TBR5 strains equally fit. Related demography-dispersal trade-offs effects have recently been observed in competing and evolving TBR1 and TBR5 yeast mats in the absence of MFs [47].

Considering the nature of microbial experiments, it will be beneficial in the future to use 3D printed materials (e.g., nylon carbon fiber) that can withstand the high temperature and pressure of an autoclave [48], as PLA loses its rigidity at temperatures between 45 *^◦^*C and 60 *^◦^*C [49]. Future work also involves repeating the TBR1-TBR5 experiments using the vertical configuration of the MF exposure device. We hypothesize that applying a vertical MF will hinder the spatial expansion of TBR5 mats perpendicular to the surface of the agar plates, but that a vertical MF will not impact TBR1 mat expansion or the growth of planktonic TBR1 or TBR5 cells. As the distance from the magnets to each agar plate is different in the vertical MF configuration, multiple MF exposure devices will be required to ensure sufficient replicates with identical MF conditions. Additionally, longer Neodymium magnets should be use in the vertical MF configuration to reduce the inhomogeneity of the 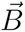 along surface of the Petri dishes. Finally, it will be imperative to elucidate the mechanisms underlying our TBR1 mat MF exposure results. Here, the magnetic properties of microtubules should be investigated in the context of yeast mat development during MF exposure. Previous studies have reported magnetic properties of microtubules [44, 50] and it has been proposed that the polarization of microtubules is responsible for the alignment of *S. cerevisiae* in the direction of an applied MF [11]. As radical pairs may play a role in microtubule reorganization, they should also be investigated as possible mechanism underlying magnetic phenomena in yeast [51]. Overall, we anticipate that our novel MF exposure device and experimental findings will advance our fundamental understanding of magnetic phenomena in fungi.

## Supporting Materials

Supporting materials includes seven figures, three tables, and eight supplemental figures.

Experimental data available at: https://data.mendeley.com/datasets/ycm2rgcfdx/1.

Exposure device design DIY is available at: https://github.com/CharleboisLab/Exposure-Design-DIY---AutoCAD-and-STL-Files.git.

## Acknowledgments

The authors thank Prof. Mark Freeman for guidance on 3D printing and for access to a Gaussmeter and Prof. Jack Tuszynski for helpful discussions.

DC was supported by funding from the Government of Canada’s NSERC Discovery Grant (RGPIN-2020-04007) and Launch Supplement (DGECR-2020-00197). EL was supported by the University of Alberta’s URI Undergraduate Researcher Stipend. This work was completed in part with resources provided by The Shack at the University of Alberta to 3D print prototypes of the magnetic field device.

## Author contributions

DC conceptualised the study. AB designed, modeled, built, and validated the magnetic field device. AB performed the yeast experiments with assistance from EL. AB visualised and analysed the results. DC and AB wrote the manuscript. DC supervised the study.

## APPENDICES

## Appendix A. Parameters

**Table A1:** Parameter values used in the COMSOL simulation of the magnetic field exposure device. The values for the N52 grade Nd_2_Fe_14_B magnets were obtained from the COMSOL material library [33].

| Parameter | Value |
| --- | --- |
| Permeability of Vacuum ($\mu_0$) | $4\pi \times 10^7 \text{ Hm}^{-1}$ |
| Relative Permeability of Air ($\mu_{r,air}$) | 1.00 |
| Relative Permeability of PLA ($\mu_{r,PLA}$) | 1.00 |
| Relative Permeability of Petri Dishes ($\mu_{r,Petri}$) | 1.00 |
| Recoil Permeability of N52 grade $\text{Nd}_2\text{Fe}_{14}\text{B}$ ($\mu_{rec,N52}$) | 1.05 |
| Remnant flux density of N52 grade $\text{Nd}_2\text{Fe}_{14}\text{B}$ ($B_r$) | 14400 G |

## Appendix B. Supplemental Tables

**Table B1:** Goodness of fit of the average area expansion rate data for TBR1-TBR5 control (no MF) and experimental (MF) group data. The model (linear, exponential, and logarithmic) with the best goodness of fit statistic are highlighted in green for TBR1 and yellow for TBR5.

| Parameters | Curve Fit | TBR1 Exposed | TBR1 Control | TBR5 Exposed | TBR5 Control |
| --- | --- | --- | --- | --- | --- |
| $SSE$ | Linear | **15.806** | **45.509** | 411.219 | 395.784 |
|  | Exponential | 435.510 | 781.309 | 1255.700 | 1150.900 |
|  | Logarithmic | 1306.600 | 1728.500 | **72.755** | **66.278** |
| $R^2$ | Linear | **0.999** | **0.997** | 0.964 | 0.960 |
|  | Exponential | 0.959 | 0.947 | 0.889 | 0.883 |
|  | Logarithmic | 0.878 | 0.883 | **0.994** | **0.993** |
| $AdjR^2$ | Linear | **0.999** | **0.997** | 0.962 | 0.958 |
|  | Exponential | 0.957 | 0.944 | 0.884 | 0.878 |
|  | Logarithmic | 0.872 | 0.876 | **0.993** | **0.993** |
| $RMSE$ | Linear | **0.829** | **1.687** | 4.228 | 4.148 |
|  | Exponential | 4.351 | 6.988 | 7.389 | 7.074 |
|  | Logarithmic | 7.537 | 10.394 | **1.779** | **1.698** |

**Table B2:** Goodness of fit of average area data for TBR1-TBR5 control (no MF) and experimental (MF) group data. The model (linear, exponential, and logarithmic) with the best goodness of fit statistic are highlighted in green for TBR1 and yellow for TBR5.

| Parameters | Curve Fit | TBR1 Exposed | TBR1 Control | TBR5 Exposed | TBR5 Control |
| --- | --- | --- | --- | --- | --- |
| $SSE$ | Linear | 405950.0 | 252690.0 | **195520.0** | **159100.0** |
|  | Exponential | **227490.0** | **192420.0** | 420520.0 | 414560.0 |
|  | Logarithmic | 3010800.0 | 1842900.0 | 2691900.0 | 2340400.0 |
| $R^2$ | Linear | 0.961 | 0.963 | **0.982** | **0.984** |
|  | Exponential | **0.978** | **0.972** | 0.962 | 0.958 |
|  | Logarithmic | 0.707 | 0.728 | 0.756 | 0.760 |
| $AdjR^2$ | Linear | 0.959 | 0.960 | **0.982** | **0.983** |
|  | Exponential | **0.977** | **0.970** | 0.960 | 0.956 |
|  | Logarithmic | 0.695 | 0.711 | 0.746 | 0.750 |
| $RMSE$ | Linear | 132.853 | 125.670 | **92.201** | **93.170** |
|  | Exponential | **99.452** | **109.665** | 135.216 | 134.255 |
|  | Logarithmic | 361.805 | 339.385 | 342.113 | 318.990 |

The following statistics were used to evaluate the goodness of fit for the data in Table B1 and Table B2. The *SSE* - sum of squares due to error:

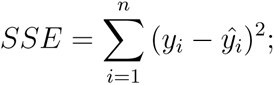

*R*^2^ - ratio between the sum of squares of the regression (*SSR*) and the total sum of squares (*SST*):

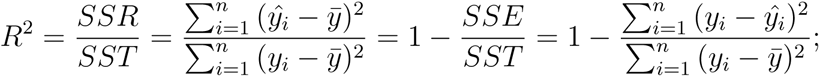

*AdjR*^2^ - degrees of freedom adjusted *R*^2^:

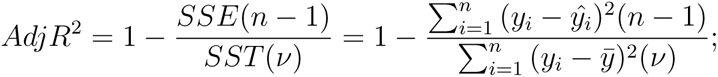

and *RMSE* - root mean squared error:

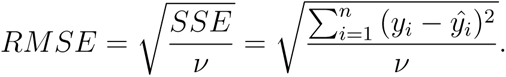

For the above equations, *y_i_* is the *i^th^* value of the variable to be predicted, *ŷ_i_* the predicted value of *y_i_*, *y̅* the mean of all values of *y_i_*, *n* the number of data points, *ν* the number of number of degrees of freedom, and (*ν* = *n − m*), where *m* is the number of fitted coefficients estimated from the data points.

## Appendix C. Supplemental Figures

**Figure C1:**
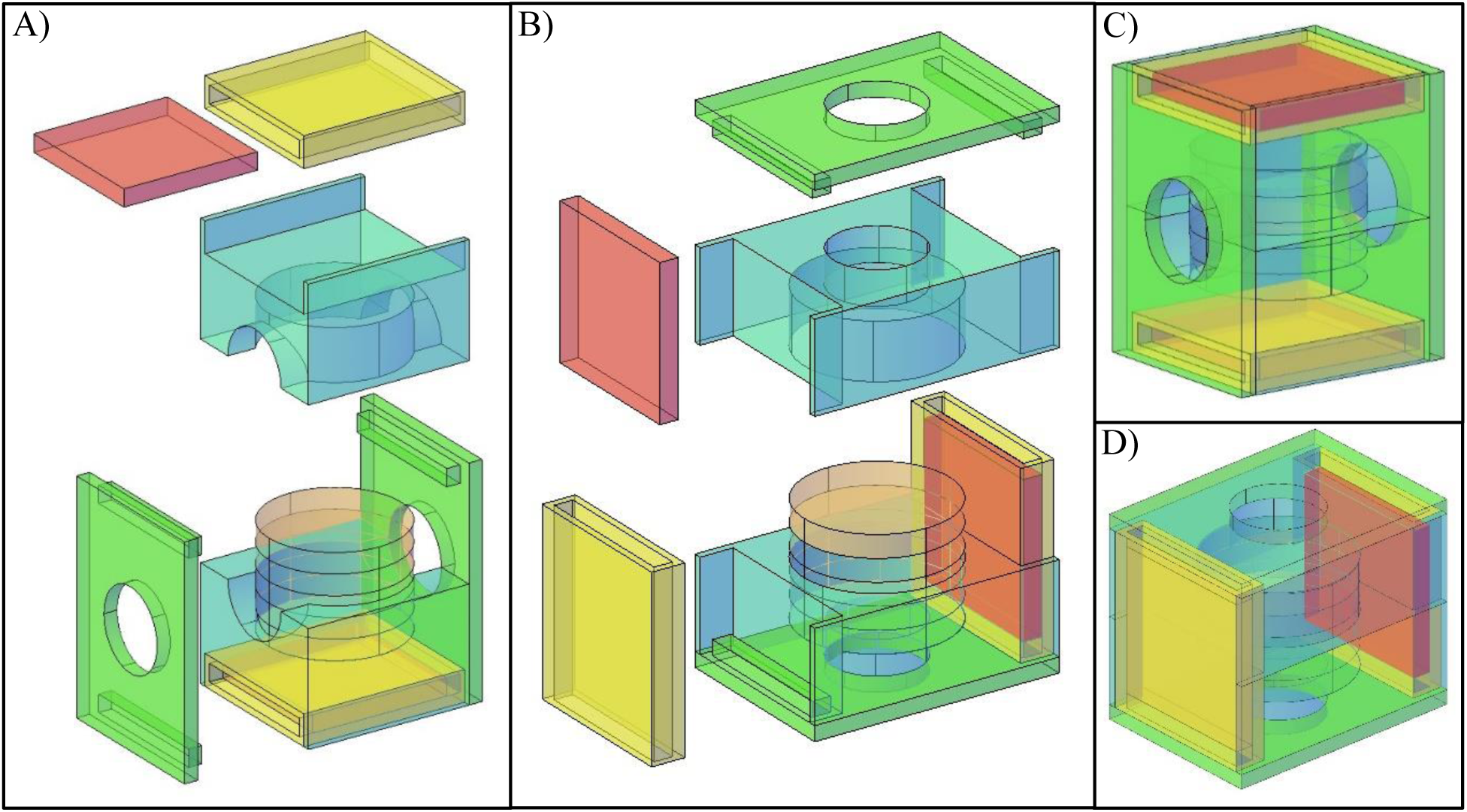
Modular design of the magnetic field exposure device. (A) AutoCAD [32] image of the disassembled vertical magnetic field (MF) configuration of the device. (B) AutoCAD image of the disassembled horizontal MF configuration of the device. (C) Assembled AutoCAD image of the vertical MF configuration of the device. (D) Assembled AutoCAD image of the horizontal MF configuration of the device. The magnets are depicted in red, magnet holders in yellow, Petri dish holders in cyan, Petri dishes in orange, and the yokes (parts that hold the device together after assembly) in green.

**Figure C2:**
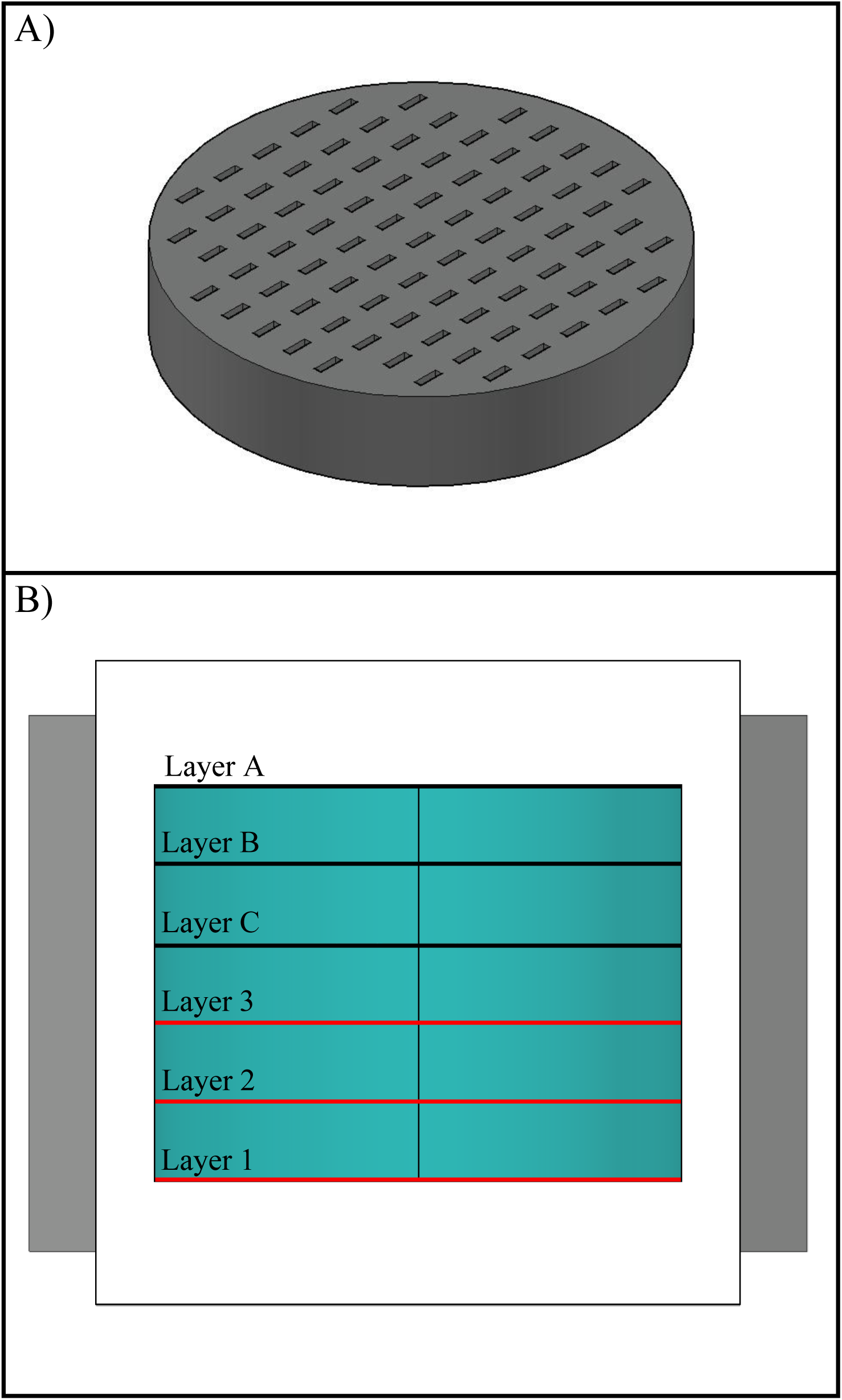
Experimental setup to map the magnetic flux density (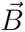). (A) AutoCAD [32] image of the cylindrical device with 83 rectangular holes used to hold the Gaussmeter probe during 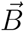 measurements. (B) Schematic of three different layers in which 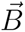 was mapped using the Gaussmeter. The grey blocks denote the permanent magnets and the Petri dishes are shown in blue. The red line indicate the layers that were evaluated in the 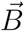 mapping.

**Figure C3:**
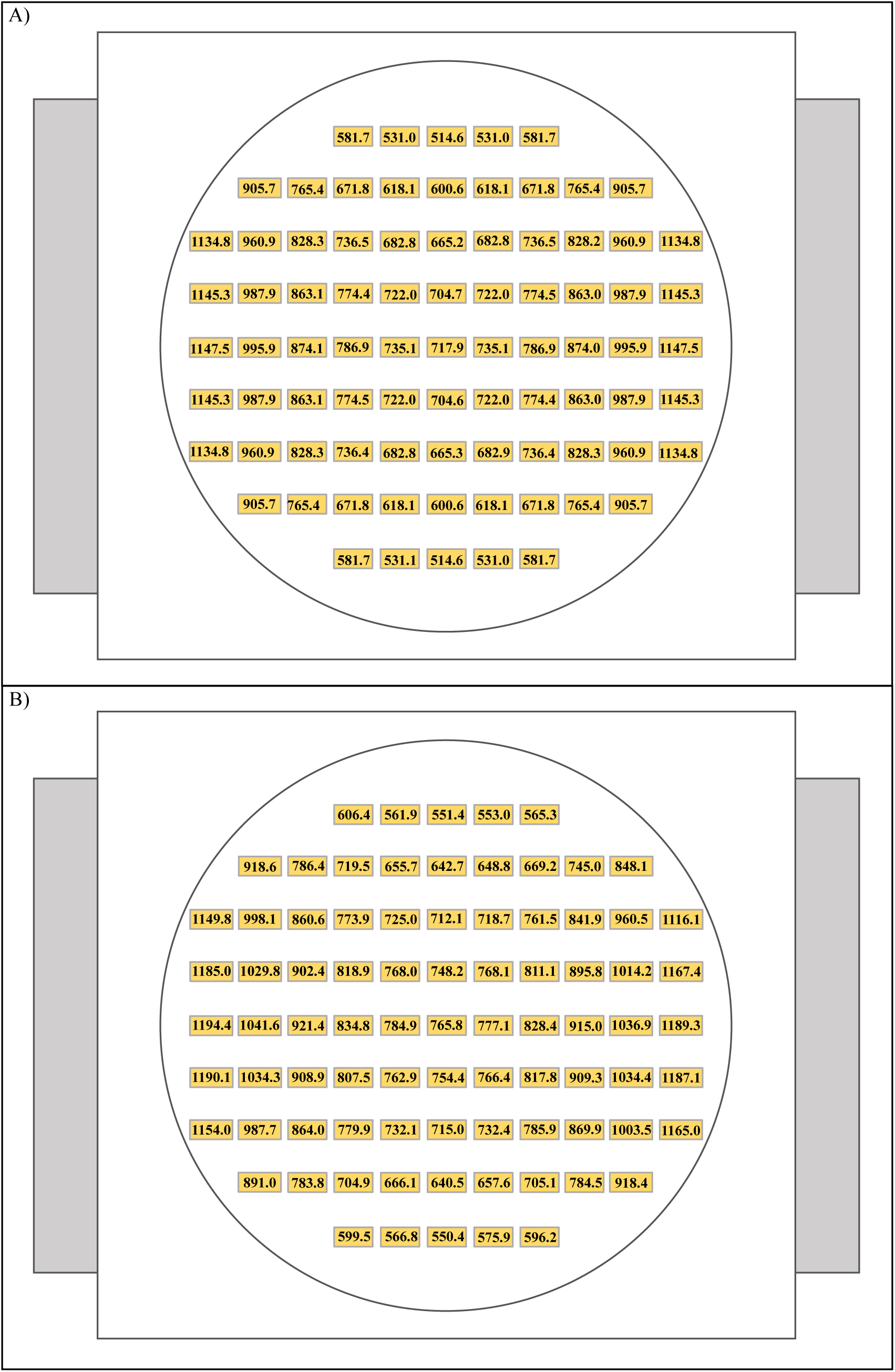
Simulated and experimentally measured values of the magnetic flux density of the horizontal configuration of the magnetic field device for layer 3. The point of view for this figure is from the top view of the middle layer (layer 3). (A) Magnetic flux density (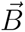) values obtained from a COMSOL [33] simulation. (B) 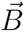 values obtained from Gaussmeter measurements. Values of 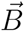 in (A) and (B) are in Gauss (*G*).

**Figure C4:**
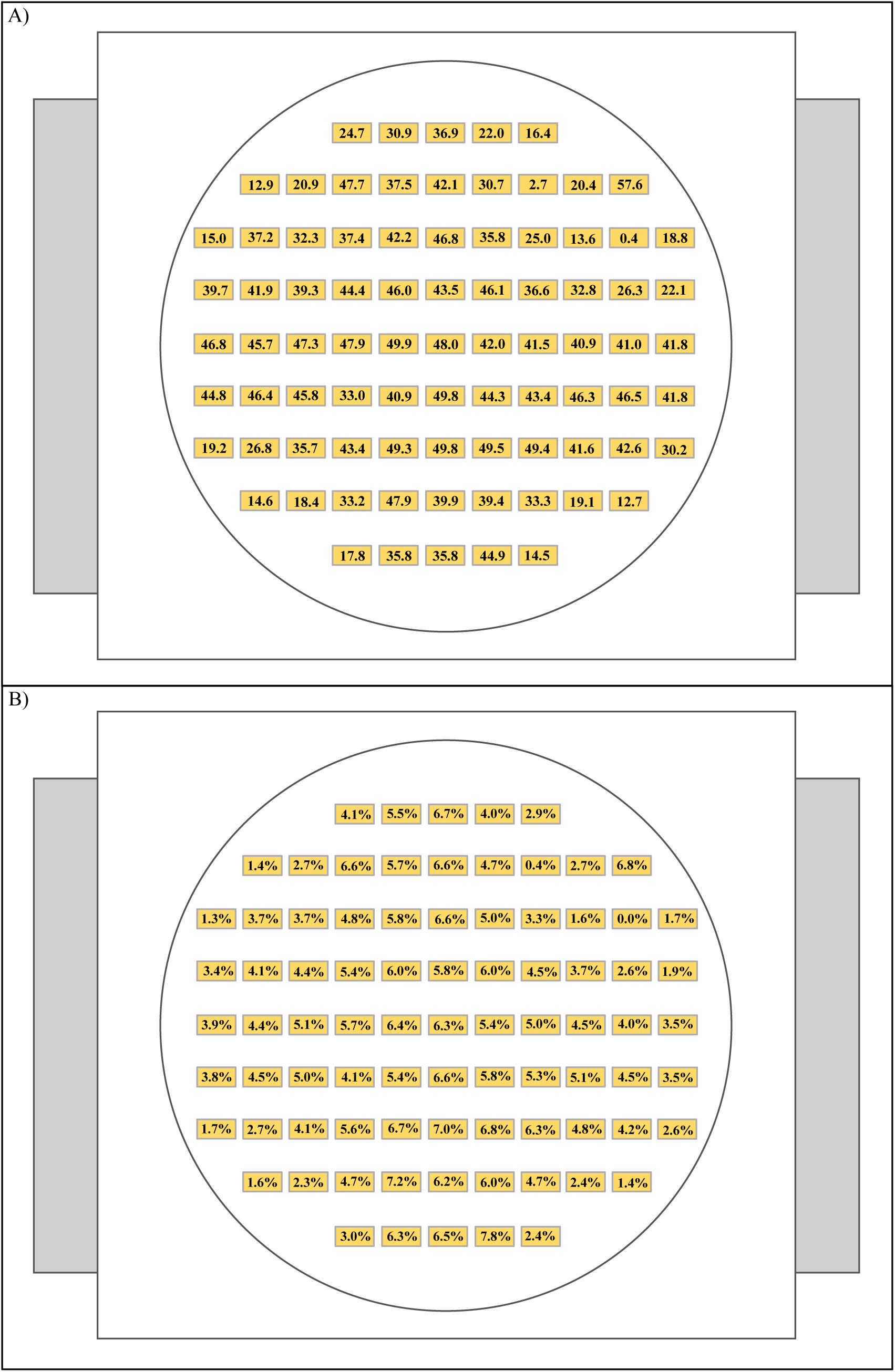
Difference between simulated and experimental magnetic flux densities of horizontal configuration of the magnetic field device for layer 3. The point of view for this figure is from the top view of the middle layer (layer 3). (A) The difference between the simulated and measured magnetic flux densities (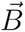). Values of 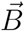 are in Gauss (*G*). (B) The difference between the simulated and experimentally measured 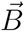 values as a percentage of the experimental values.

**Figure C5:**
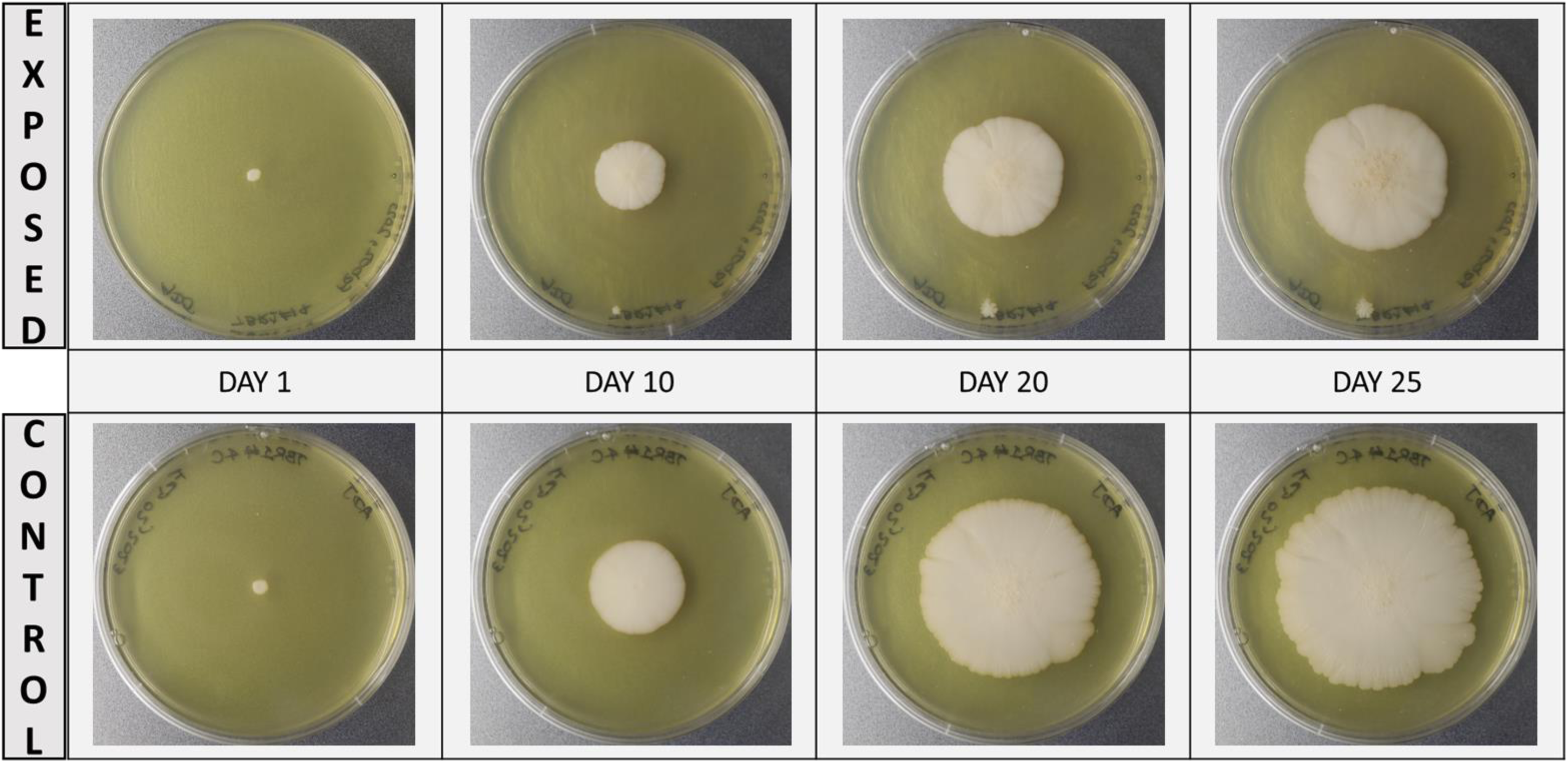
Representative images of the development of TBR1 yeast mats for (Top) the exposed condition (magnetic field) and for (Bottom) the control condition (no magnetic field).

**Figure C6:**
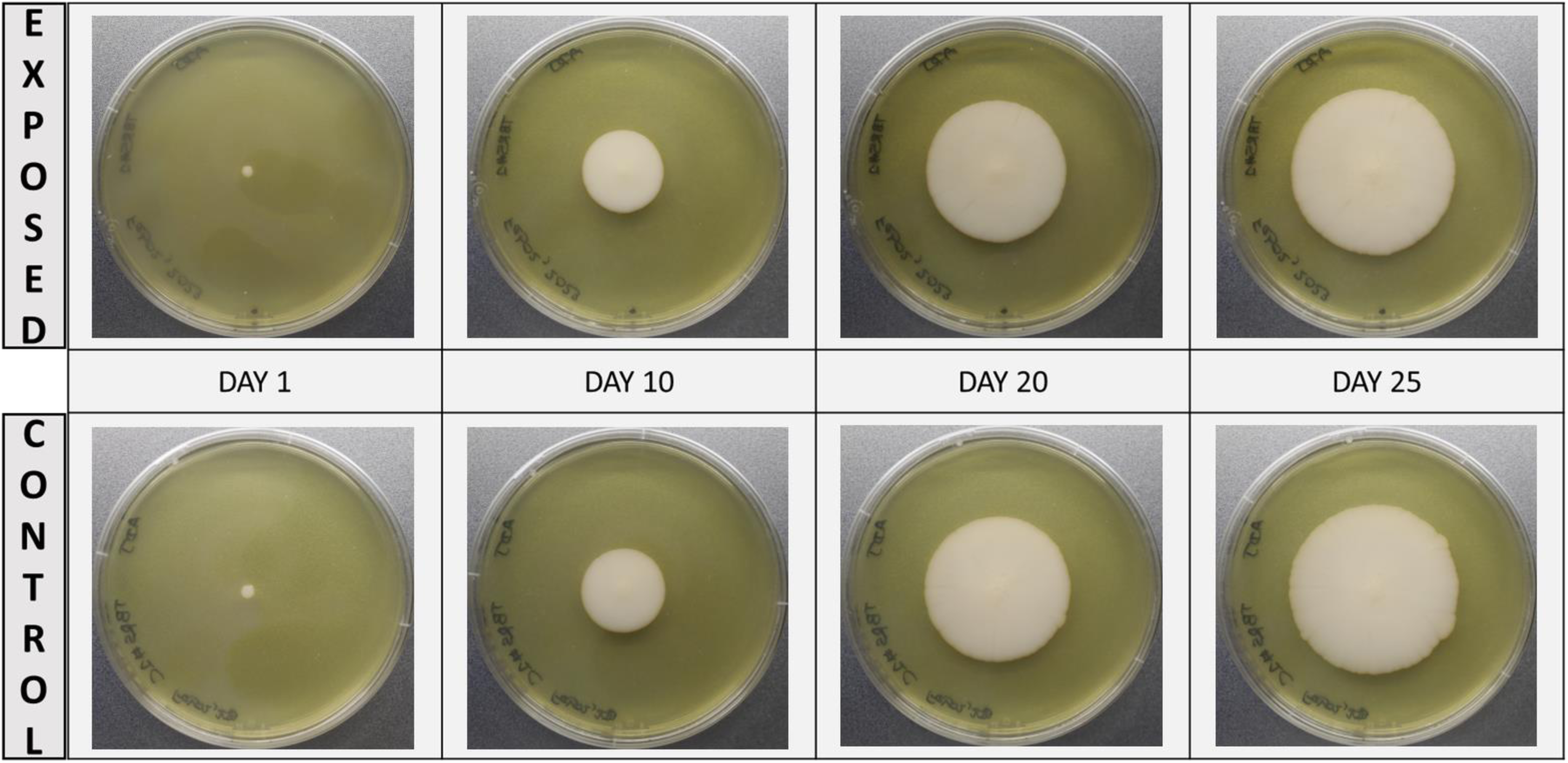
Representative images of the development of TBR5 yeast mats for (Top) the exposed condition (magnetic field) and for (Bottom) the control condition (no magnetic field).

**Figure C7:**
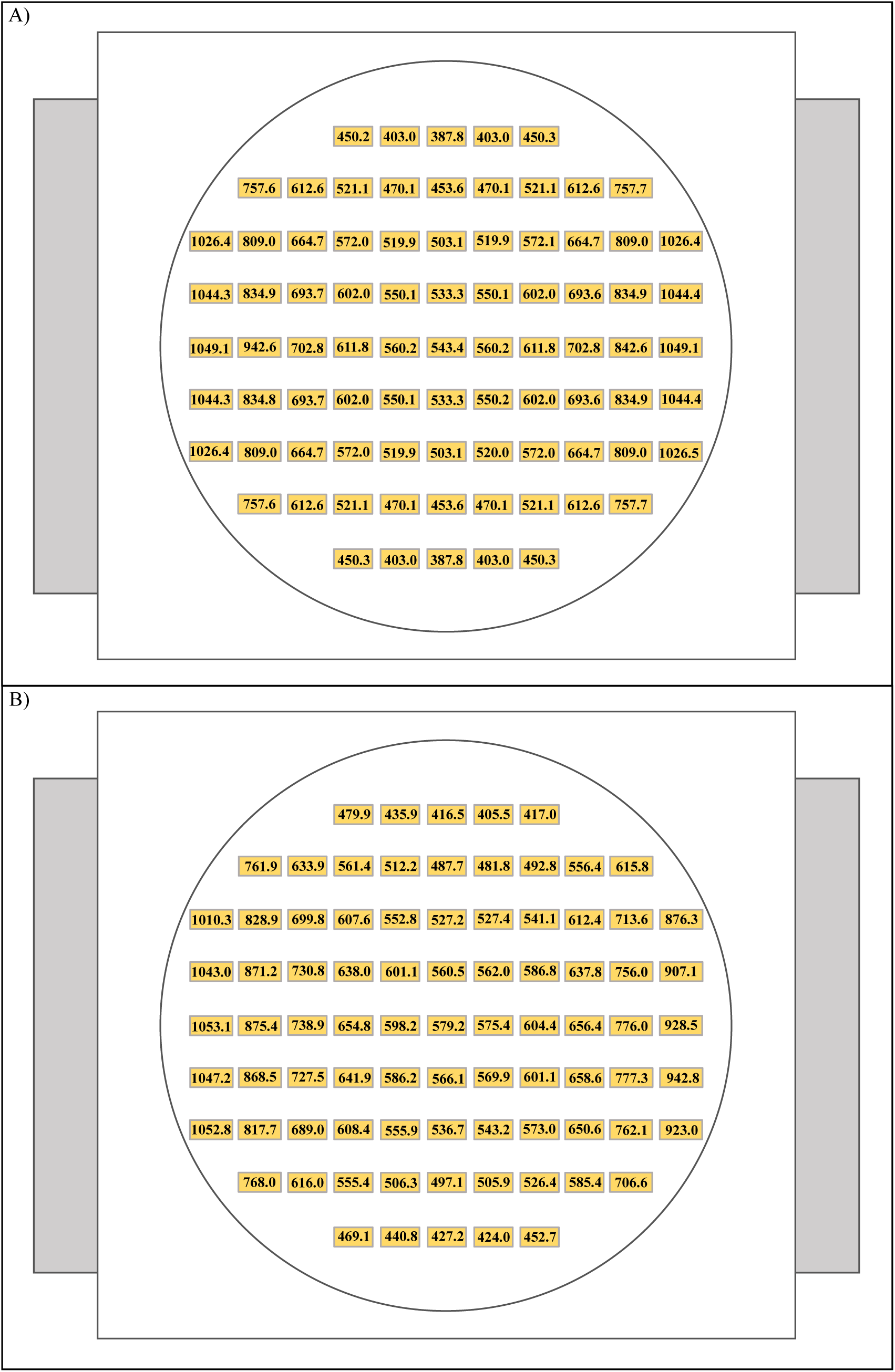
Simulated and experimentally measured values of the magnetic flux density of the horizontal configuration of the magnetic field device for layer 1. The point of view for this figure is from the top view of the layer 1. (A) Magnetic flux density (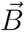) values obtained from a COMSOL [33] simulation. (B) 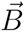 values obtained from Gaussmeter measurements. Values of 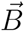 in (A) and (B) are in Gauss (*G*).

**Figure C8:**
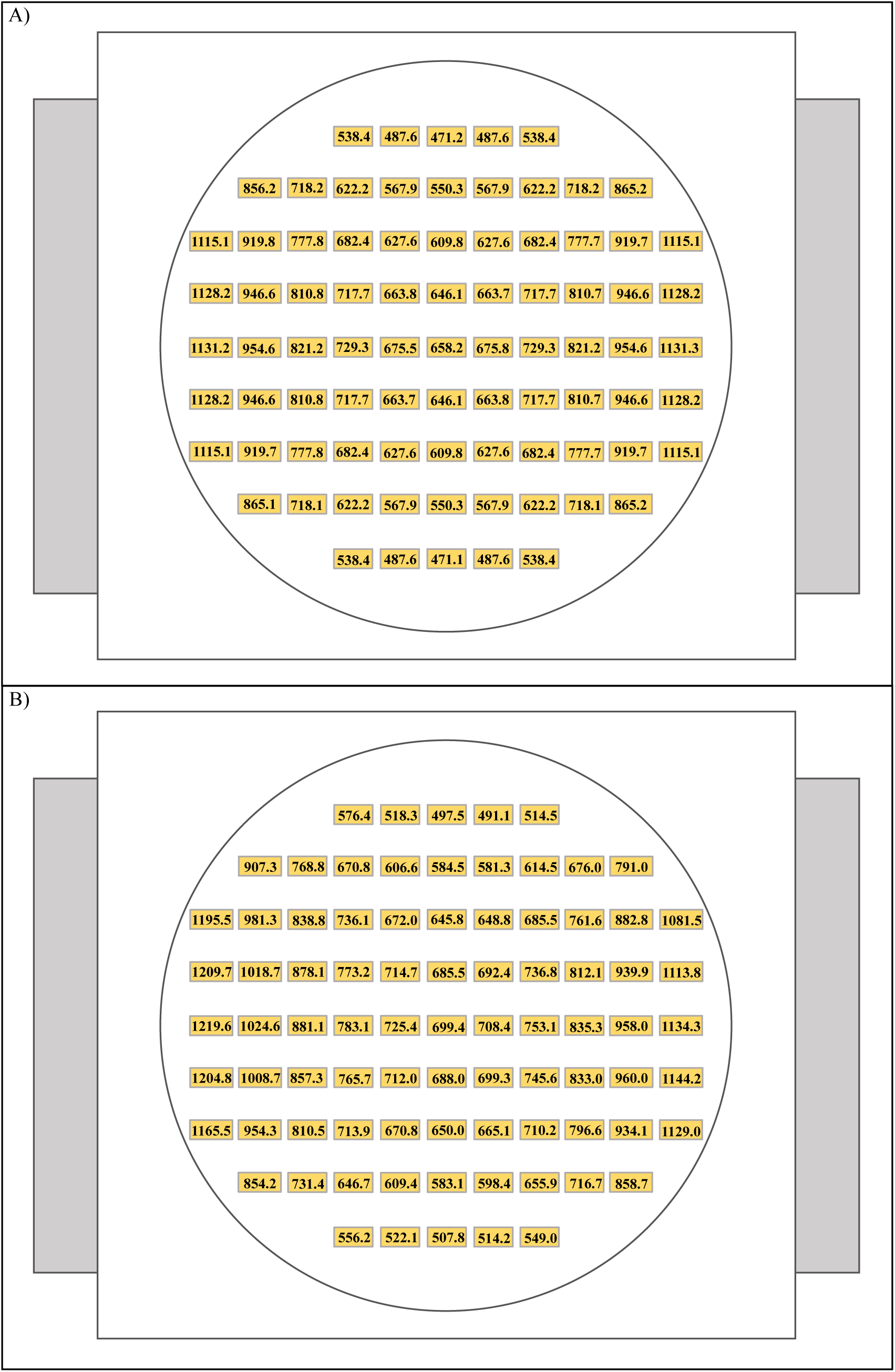
Simulated and experimentally measured values of the magnetic flux density of the horizontal configuration of the magnetic field device for layer 2. The point of view for this figure is from the top view of the layer 2. (A) Magnetic flux density (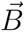) values obtained from a COMSOL [33] simulation. (B) 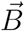 values obtained from Gaussmeter measurements. Values of 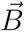 in (A) and (B) are in Gauss (*G*).

